# Single-shot rAAV5-based Vaccine Provides Long-term Protective Immunity against SARS-CoV-2 and Its Variants

**DOI:** 10.1101/2021.08.23.456471

**Authors:** Guochao Liao, Xingxing Fan, Hungyan Lau, Zhongqiu Liu, Chinyu Li, Zeping Xu, Yu Zhang, Xiaoxiao Qi, Dan Li, Qing Zhu, Liqing Chen, Hua Zhou, Sisi Zhu, Bixia Ke, Hudan Pan, Zhe Cong, Yongchao Li, Qian Feng, Qi Lv, Jiangning Liu, Dan Liang, An’an Li, Wenshan Hong, Yebo Li, Linlin Bao, Feng Zhou, Hongbin Gao, Shi Liang, Bihong Huang, Miaoli Wu, Chuan Qin, Changwen Ke, Liang Liu

## Abstract

The COVID-19 pandemic and the SARS-CoV-2 with its variants have posed unprecedented challenges worldwide. Existing vaccines have limited effectiveness against the SARS-CoV-2 variants. Therefore, novel vaccines to match current mutated viral lineages with long-term protective immunity are urgently in demand. In the current study, we for the first time designed a recombinant Adeno-Associated Virus 5 (rAAV5)-based vaccine named as rAAV-COVID-19 vaccine (Covacinplus) by using RBD-plus of spike protein with both the single-stranded and the self-complementary AAV5 delivering vectors (ssAAV5 and scAAAV5), which provides excellent protection from SARS-CoV-2 infection. A single dose vaccination induced the strong immune response against SARS-CoV-2. The induced neutralizing antibodies (NAs) titers were maintained at a high peak level of over 1:1024 even after more than one year of injection and accompanied with functional T-cells responses in mice. Importantly, both ssAAV- and scAAV-based RBD-plus vaccines exhibited high levels of serum NAs against current circulating variants including variants Alpha, Beta, Gamma and Delta. SARS-CoV-2 virus challenge test showed that ssAAV5-RBD-plus vaccine protected both young and old age mice from SARS-CoV-2 infection in the upper and the lower respiratory tracts. Moreover, whole genome sequencing demonstrated that AAV vector DNA sequences were not found in the genome of the vaccinated mice after one year vaccination, demonstrating excellent safety of the vaccine. Taken together, this study suggests that rAAV5-based vaccine is powerful against SARS-CoV-2 and its variants with long-term protective immunity and excellent safety, which has great potential for development into prophylactic vaccination in human to end this global pandemic.

## Main text

Severe acute respiratory syndrome coronavirus-2 (SARS-CoV-2) has caused more than 209 million people infections and 4.4 million deaths as of Aug 21, 2021; while the global pandemic of COVID-19 has provided a large number of opportunities for mutation. Especially, the circulating variants of SARS-CoV-2 of Delta resulted in accelerated worldwide spreading of the virus. Various studies showed that the Delta variant is more infectious and dangerous, reducing the immune efficacy and protection of existing vaccines, inducing immune escape and leading to re-infection in people who have recovered from the wild-type SARS-CoV-2^1,2^. Currently, there are nearly 100 vaccines in clinical trials and 200 pre-clinical vaccine candidates^3–5^, however, few of these have shown significant protective efficacy against both SARS-CoV-2 and its circulating variants in the long term, the Delta variant in particular^1,6^. Therefore, an effective vaccine against wild-type SARS-CoV-2 and its variants is urgent in demand.

Adeno-Associated Virus (AAV) has been widely used for gene therapy due to its safety and long-term characteristics, with high efficacy and low immunogenicity^7–9^, which highly addressed the requirements of ideal vaccine vectors. The prevalence of pre-existing NAs (PNAs) against AAV vectors is directly related to vaccine efficacy^10^; while AAV type 5 (AAV5) with low PNAs (3.2%)^11,12^ and high affinity to human airway epithelia^13^ possesses a great potential to be developed as the most optimal vaccine vector. Moreover, the safety of AAV5 vector in human has been well studied and evaluated^14^. Although currently recombinant engineering AAV vector has been used for effective gene delivery^15,16^, whether it is compatible to human still requires clinical trial to prove. Therefore, we utilized AAV5 delivery vehicle to produce a broad and long-term protective vaccine against SARS-CoV-2 and its variants.

In order to systematically screen the most effective SARS-CoV-2 spike (S) protein antigen for inducing immune response, we first designed five single-stranded DNA recombinant AAV5-based (ssAAV5) vaccines: ssAAV5-RBD (encodes the residues 319-541 RBD of S protein), ssAAV5-RBD-plus (encodes the residues 331-583 RBD-plus of S protein), ssAAV5-S1 (encodes the residues 14-685 of S1), ssAAV5-S (encodes the residues 14-1213 of extracellular domain of S protein), and ssAAV5-NTD (encodes the residues 14-305 of NTD) (Fig.1A). Given sequence conservation of different domains on S protein, we extended the length of antigen in the RBD-plus domain to compare with the mere RBD domain. The immunogenicity of designed vaccines was assessed in BALB/c mice. Animals were immunized by intramuscular (i.m) inoculation with 1 × 10^11^ genome copies (GCs) of each vaccine or parallel control (AAV5-GFP), respectively (1 × 10^12^ GCs/mL, 100 μL). The antigen-specific humoral immune responses of immunoglobulin G (IgG) (Fig.1B) and NA titers in wild-type SARS-CoV-2 infected Vero E6 cells were evaluated on day 40 and continued until day 130 after single dose immunization (Fig.1C). Among these groups, ssAAV5-RBD-plus elicited the highest specific IgG and NA titers, which indicates that RBD-plus domain with residues 331-583 was able to elicit the strongest immune response against wild-type SARS-CoV-2 in comparison with other antigens. Meanwhile, T-cell responses after vaccination were evaluated. After 130 days of ssAAV5-RBD-plus vaccine immunization, populations of the memory CD4^+^ and CD8^+^ T cells were found to be significantly increased (Fig. 2A and 2B) along with significant upregulation of cytokine IFN-γ secretion from CD8^+^ compared to the controlled animals (Fig. 2C). These results indicate that ssAAV5-based RBD-plus vaccine could effectively induce both robust systemic humoral and cell-mediated immune responses.

**Fig. 1.**
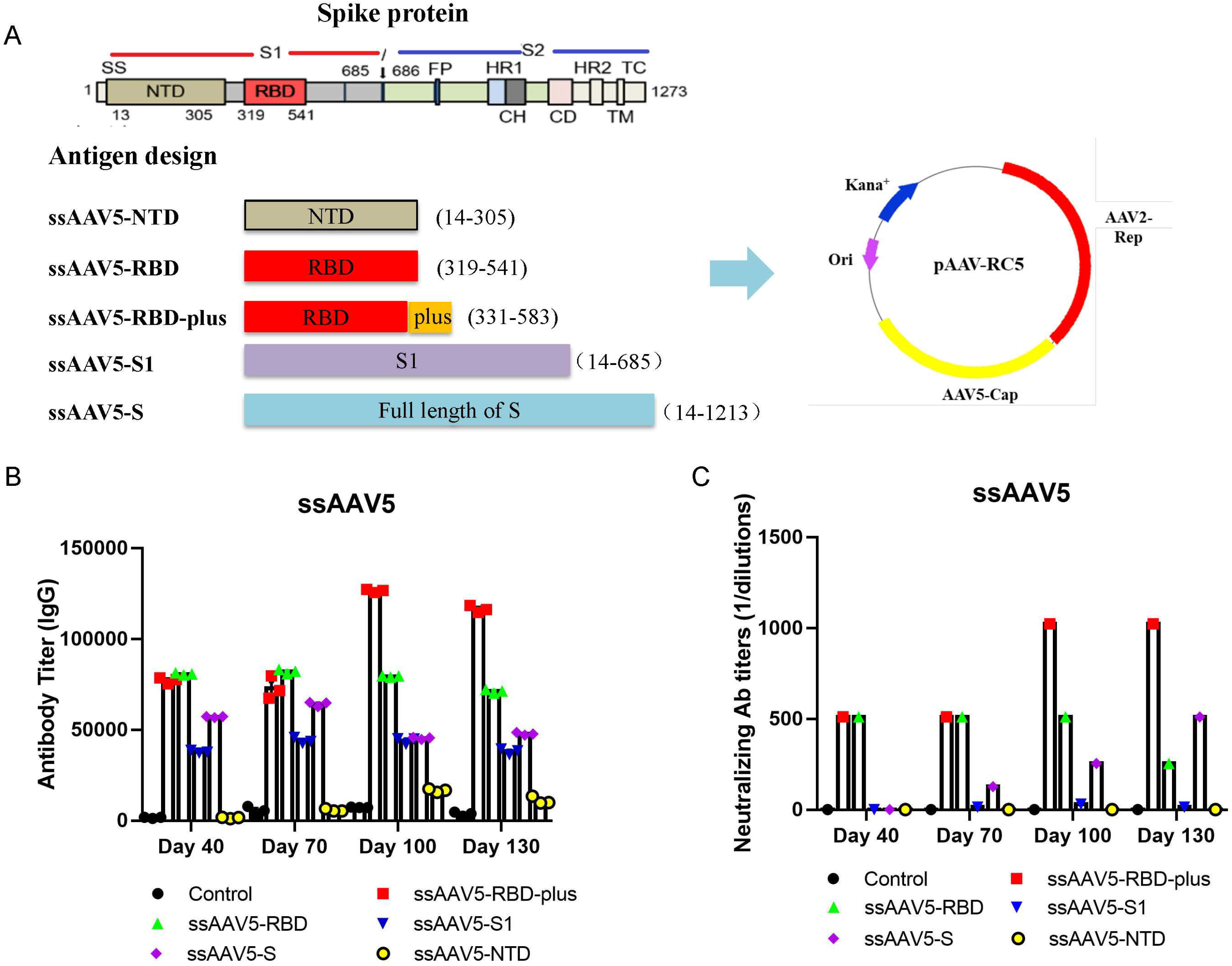
Construction of the recombinant AAV5-based vaccines and their immunogenicity; (A) Design of five recombinant ssAAV5-based vaccines; (B) IgG and (C) Wild-type virus-specific NAs were evaluated on 40 days and continued until 130 days after a single dose immunization. ssAAV5-RBD-plus vaccine induced the strongest immune responses in animals.

**Fig. 2.**
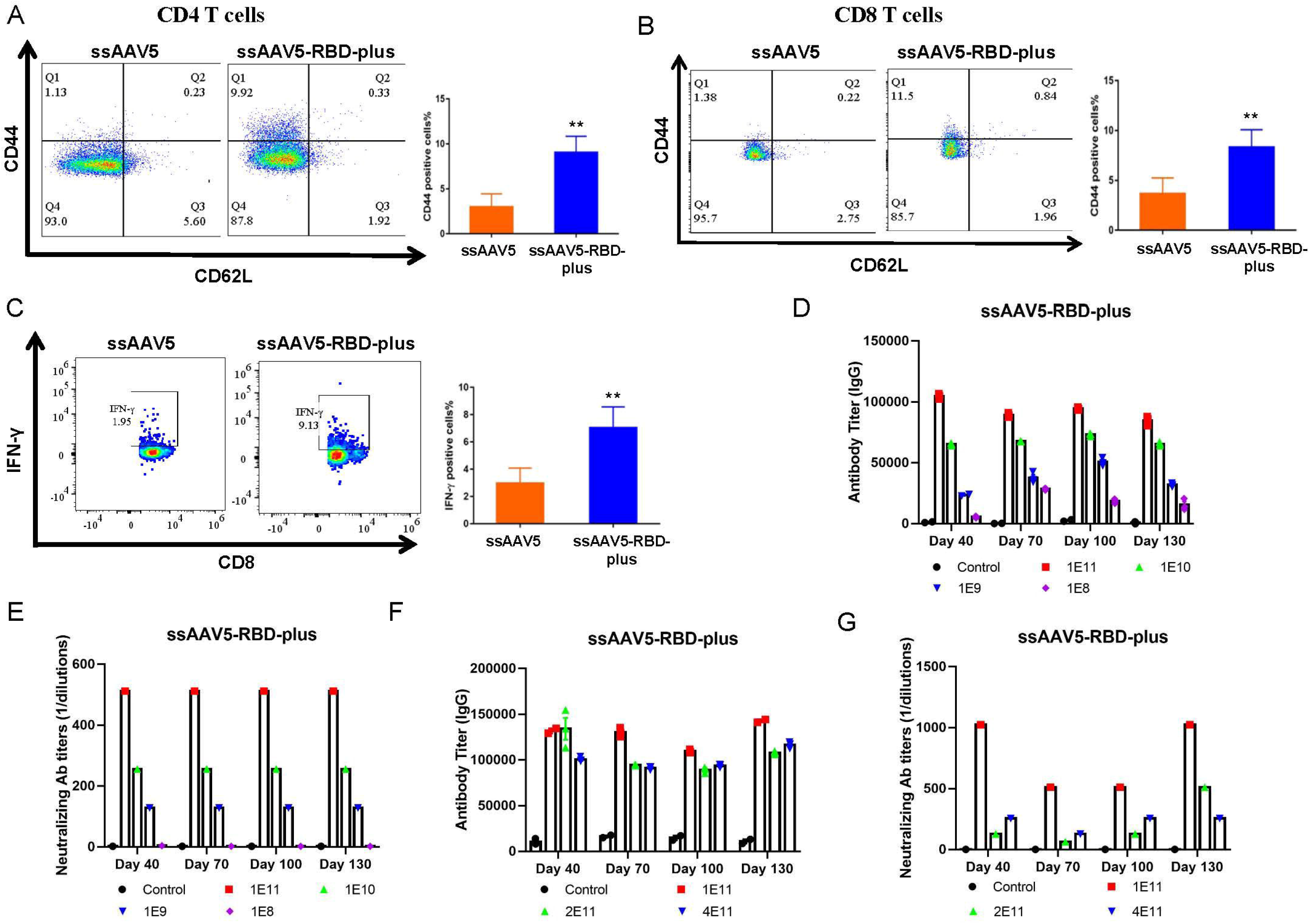
Determination of T cell immune responses and the most effective dosage of vaccine. (A) After 130 days of immunization with ssAAV5-RBD-plus vaccine, the populations of memory CD4^+^ T and CD8^+^ cells (B) were significantly increased; (C) Secretion of IFN-γ from CD8^+^ was significantly upregulated; Among dosages of 1×10^11^, 1×10^10^1, 1×10^9^, 1×10^8^ GCs, the amount of IgG (D) and NAs (E) indicates that 1×10^11^ GCs possessed the strongest immunogenicity; Among dosages of 1×10^11^, 2×10^11^ and 4×10^11^ GCs, the amount of IgG (F) and NAs (G) 1×10^11^ GCs possessed the strongest immunogenicity. Values represent mean ± SEM, significant difference versus control group, **, *P*< 0.01.

Effective optimal dosage is a critical factor for a vaccine. To find out the optimal dosages of ssAAV5-RBD-plus vaccine, we first vaccinated mice by i.m at dosages of 1×10^8^, 1×10^9^, 1×10^10^, and 1×10^11^ GCs, respectively. The results showed that the titers of IgG and NA were elevated with increasing of dosages while the animals vaccinated with 1×10^11^ GCs showed the strongest immunogenicity (Fig. 2D and 2E). To further clarify whether 1×10^11^ GCs was the most effective dosage, higher dosages with 2×10^11^ and 4×10^11^GCs were vaccinated to mice. As shown in Fig. 2F and 2G, animals vaccinated with 1×10^11^ GCs obtained the strongest immune responses. Based on these results, 1×10^11^ GCs in the ssAAV5-based vaccine were determined as the optimal dose for further studies.

Then, SARS-CoV-2 virus challenge test was applied to determine the protective effect of ssAAV5-RBD-plus vaccine on 6-week old (Young) and 36-week old (Old) BALB/c mice by a single dose of 6×10^10^ GCs, respectively, with two routes of vaccine administration, namely i.m and intranasal routs. After 40 days of vaccination, mice were challenged intranasally with 3.6Log PFU of SARS-CoV-2 (HRB26M strain)^17^. The results showed that both of the administration routes of ssAAV5-RBD-plus vaccine significantly increased the NAs both in young and old BALB/c mice (Fig. 3A and 3B). After 3 and 5 days of viral attack, the titers of virus were measured both in the lungs and nose. As shown in Fig. 3C and 3D, the number of viruses dropped nearly 4.0 Log copies/g in lung tissues, which indicates that ssAAV5-RBD-plus vaccine provided almost full protection for lungs, suggesting that 6×10^10^ GCs is an effective dosage of ssAAV5-RBD-plus vaccine to induce immune protection against SARS-CoV-2. The titers of infectious virus in nose were higher than that in lungs (Fig. 3E and 3F). Together, a single dose of ssAAV5-RBD-plus vaccine exhibited strong protection against wild-type SARS-CoV-2 in contrast to other vaccines that require multiple vaccinations^1,18^.

**Fig. 3.**
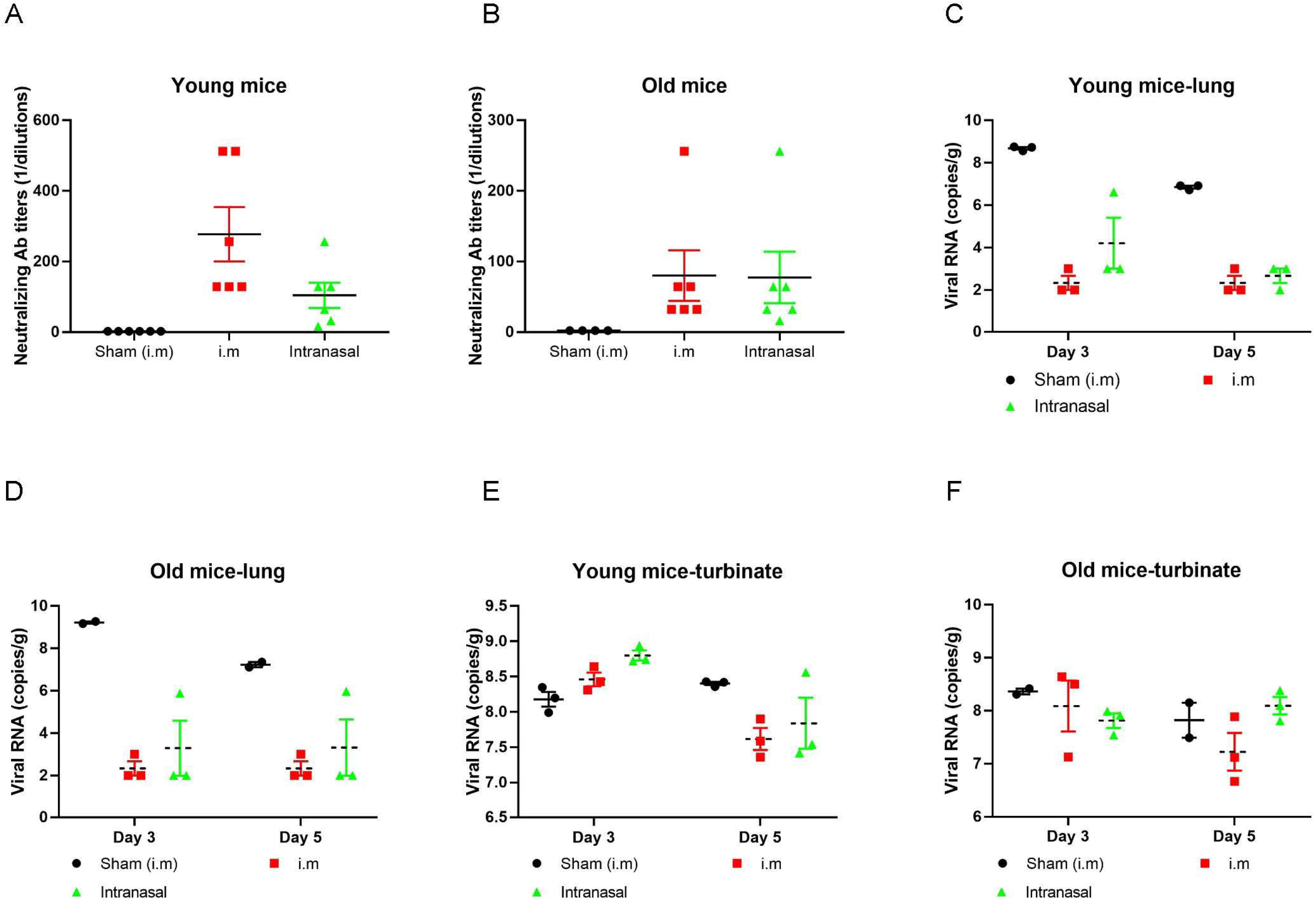
SARS-CoV-2 virus challenge test on young and old mice. (A) and (B) ssAAV5-RBD-plus significantly increased the NAs in both young and old BALB/c mice; (C) and (D) RBD-plus provided excellent protection for lung, showing the number of viruses dropped by more than 4.0 Log copies/g of in lung tissues of the vaccinated young and old mice; (E) and (F) After 3 and 5 days of virus inoculation, the titers of infectious virus on nose dropped nearly 1.0 Log copies/g in both young and old mice.

Multiple studies have demonstrated two major requirements for vaccines to efficiently provide strong, long-lasting neutralization titers and broad protection against SARS-CoV-2 variants^19–21^. Mutations in the S protein in circulating variants of SARS-CoV-2 reduce the efficacy of antibody neutralization induced by previous vaccination^22^. Especially, mutation in RBD domain can impede the binding of NAs with the virus, which results in low efficiency or even ineffectiveness of vaccines targeting wild-type SARS-CoV-2. Moreover, the neutralizing activity declines dramatically or even loses within couple of months after infection^23^. Therefore, we investigated the long-term protective effect of ssAAV5-RBD-plus vaccine after 1 year of immunization on various variants. Sera samples of mice were collected on day 376 after a single-shot immunization and humoral immunity against SARS-CoV-2 was evaluated by ELISA. As shown in Fig. 4A, ssAAV5-RBD-plus vaccine induced high level of RBD-specific IgG in three different dosages. Meanwhile, ssAAV5-RBD-plus vaccine produced specifically prolonged and potent humoral responses, including the primary (IgM) (Fig. 4B) and secondary antibodies (subtypes of IgG) (Fig. 4C-F). We collected peripheral blood mononuclear cells (PBMCs) from mice after 376 days of vaccination to evaluate T cellular responses. Compared to the controlled animals, secretion of cytokines IFN-γ and IL-4 from CD4^+^ in ssAAV5-RBD-plus group increased by 4.1 and 6.5 times, respectively (Fig. 4G and 4H). These results indicated that ssAAV5-RBD-plus could induce long-term specific T cell responses after a signal short of vaccination.

**Fig. 4.**
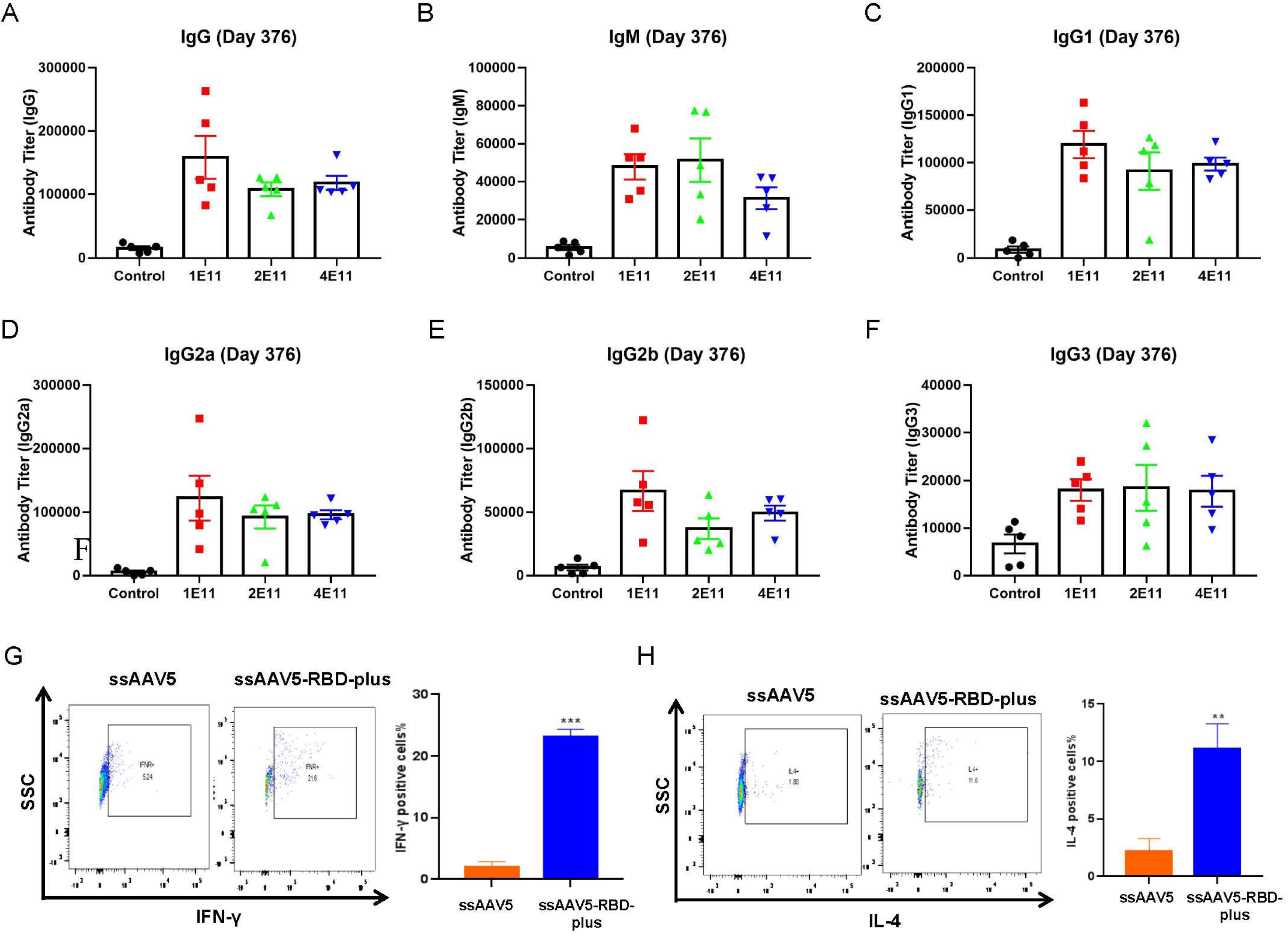
Protective effect of ssAAV5-RBD-plus vaccine after 1 year immunization on various variants. ssAAV5-RBD-plus vaccine induced high level of RBD-specific IgG (A), and specifically prolonged and potent humoral responses, including IgM (B), IgG1(C), IgG2a (D), IgG2b (E) and IgG3 (F); Secretion of IFN-γ (G) and IL-4 (H) from CD4^+^ was significantly upregulated. Values represent mean ± SEM, significant difference versus control group, **, P< 0.01, ***, P< 0.001.

We then examined the NA titers in sera from the vaccinated mice against various SARS-CoV-2 variants. In wild-type, variants Alpha and Beta, injection of ssAAV5-RBD-plus vaccine resulted in 1:1024 NAs at single dose of 1 × 10^11^ GCs, which indicates that single shot of ssAAV5-RBD-plus vaccination is capable of matching the current circulating variants (Fig. 5A). Meanwhile, the vaccination did not result in body weight loss during 376 days of experiment (Fig. 5B). Whole genome sequencing of tissues was performed in mice after one year of vaccination, and vector DNA sequences were not found in mouse genome. Collectively, the results of our study indicate that a single dose of AAV5-based RBD-plus vaccination is able to induce durable immune responses in mice and exhibits high levels of NAs against SARS-CoV-2 variants Alpha and Beta with excellent safety.

**Fig. 5.**
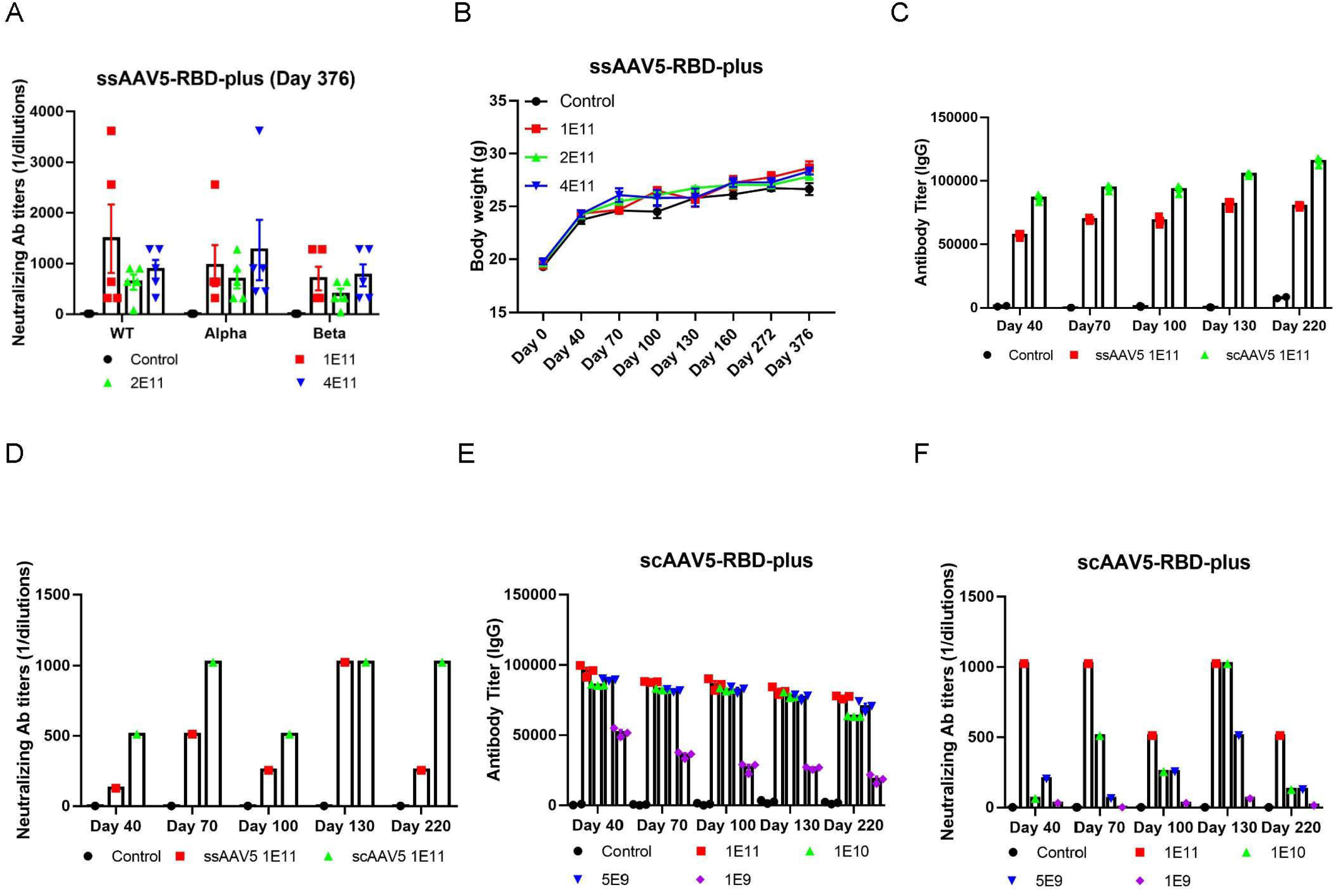
Durable immune responses and broad protection against SARS-COV-2 induced by ssAAV5-RBD-plus vaccine in mice over one-year inoculation. (A) NA titers of Wild-type (WT), Alpha and Beta SARS-COV-2; (B) Body weights of mice were increasing constantly. Immune antigenicity of ssAAV5-based and scAAV5-based vaccines in BALB/c mice; (C) IgG and (D) Wild-type virus-specific NAs were evaluated on 40 days; (E) Among all dosages of scAAV5-RBD-plus vaccines (1×10^11^, 1×10^10^, 5×10^9^ and 1×10^9^ GCs), 1×10^11^ GCs scAAV5 RBD-plus vaccine possessed the strongest IgG and NAs (F).

The efficiency of AAV delivery vectors depends on the conversion of single-stranded DNA (ssDNA) genome into double-stranded DNA (dsDNA). After ITR modification, self-complementary AAV vectors (scAAV) could bypass the rate-limiting step of second-strand synthesis and enhance the efficiency of vaccines. Therefore, we further designed a scAAV5-based vaccine encoding the residues 331-583 RBD-plus of wild-type SARS-CoV-2 and evaluated the immunogenicity of scAAV5-based RDB-plus domain vaccine in BALB/c mice. The antigen-specific humoral immune responses of IgG (Fig.5C) and virus-specific NAs (Fig.5D) were evaluated on day 40 and continued until day 220 after single dose immunization. Compared to ssAAV5-based vaccine, scAAV5-based vaccine elicited higher titers of specific IgG and NA, which indicates that scAAV5-based vaccine was able to elicit a stronger immune response against SARS-CoV-2. To verify the optimal dosages of scAAV-RBD-plus vaccine, we then vaccinated mice by i.m at dosages of 1×10^9^, 5×10^9^, 1×10^10^ and 1×10^11^ GCs, respectively. The results showed that the titers of IgG and NA were elevated with increasing of dosages while the animals vaccinated with 1×10^11^ GCs showed the strongest immunogenicity (Fig. 5E and 5F).

To further corroborate our findings, we tested the immune antigenicity of scAAV-RBD-plus vaccine in Wistar rats. The sera of the vaccinated rats were collected on days 40, 70, 100, 130, 220 and 334 after vaccine inoculation to evaluate the influences of humoral immunity and the level of NA. Results showed that scAAV vaccine was able to continuously induce production of IgG and the wild-type virus-specific NA during the 334-day experiment (Fig. 6A and 6B). After that, the sera from all vaccinated rats on day 334 were collected and the NA titers against SARS-CoV-2 variants were determined. Similar to the effect on wild-type virus, a single dose of scAAV-RBD-plus vaccine exhibited high level of serum NAs against the major current circulating variants: Alpha, Beta, Gamma and Delta variants (Fig. 6C). During the entire experimental processes, all the vaccinated rats showed no adverse responses along with continued increase in body weight (Fig.6D). These results indicated that the scAAV5-based RBD-plus vaccine (Covaccinplus) could provide a broad and long-term immune protection against SARS-CoV-2, especially for the major circulating variants such as Delta.

**Fig. 6.**
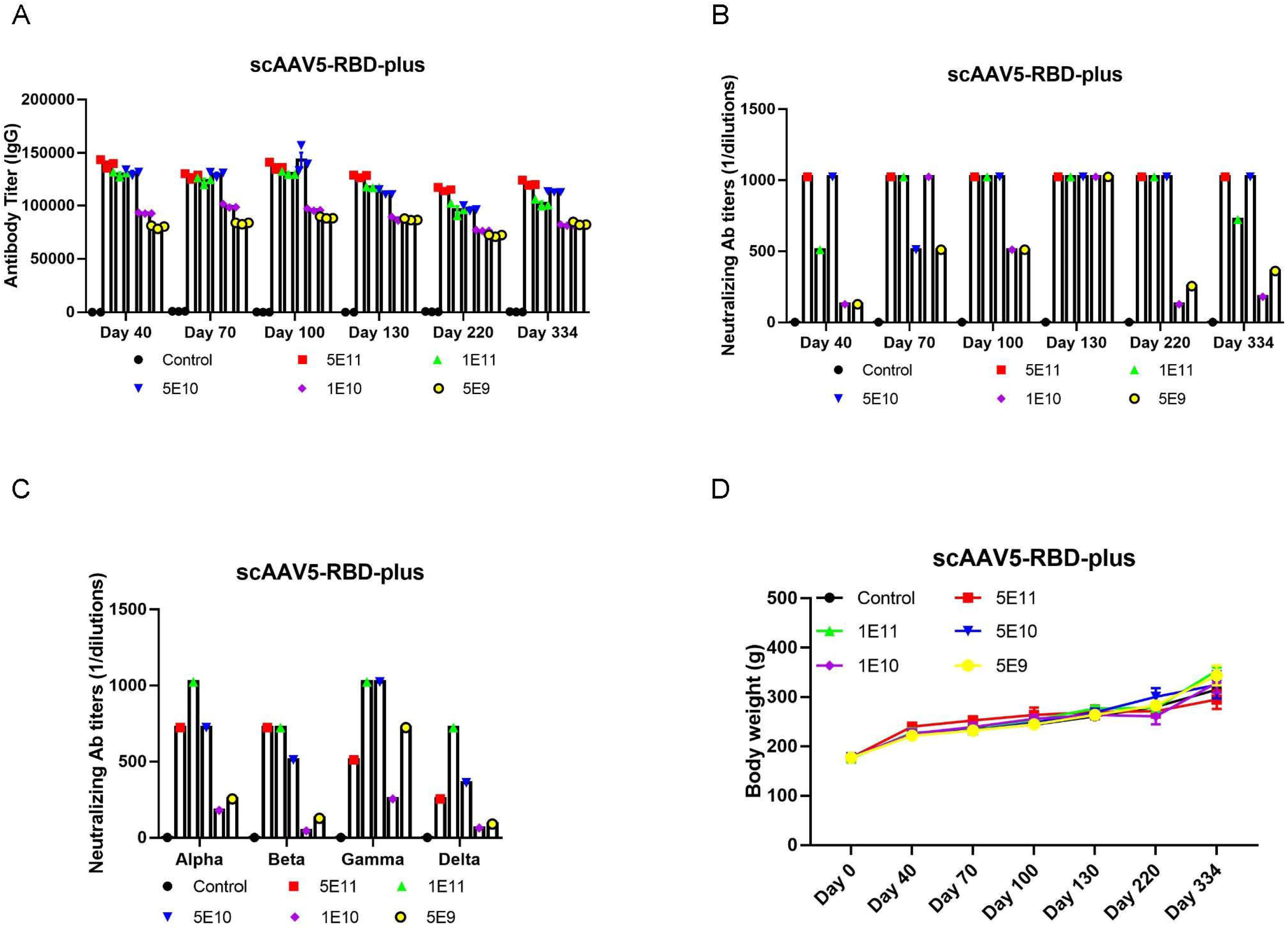
Immune antigenicity of scAAV5-RBD-plus vaccine in Wistar rats. (A) scAAV5-RBD-plus vaccine strongly induced the IgG antibody production in rats; (B) NAs against wild-type SARS-COV-2 remained at the highest level; (C) scAAV5-RBD-plus vaccine exhibited high titer of NAs against variants Alpha, Beta, Gamma and Delta SARS-COV-2; (D) Body weights of rats were increasing constantly.

In summary, both the single-stranded and the self-complementary rAAV5-based RBD-plus vaccines (Covaccinplus) represent safe and promising candidates for preventive intervention against SARS-COV-2; while our newly established recombinant rAAV5-delivery vehicle would provide a novel platform for the rational design of several new recombinant virus vector-based vaccines. These vaccines in the current study possess three unique advantages to combat the pandemic. Firstly, it induces strong and long-term immunogenicity and maintains high titers of NAs with live wide-type SARS-CoV-2 even after one year vaccination, without the need of a booster dose after primary vaccination in contrast to other vaccines^1,18^. This RBD-plus vaccine also exhibited high level of long-lasting NAs with a range of 1:500-1:1024 responding to live SARS-CoV-2 variants including Alpha, Beta and Gamma, at the optimal dosage with 1×10^11^ GCs, especially to Delta variant which is markedly resistant to the currently available vaccines. Therefore, the rAAV-based vaccine represents a powerful candidate to battle the COVID-19 pandemic^15^. Secondly, as a non-pathogenic and non-enveloped virus, rAAV5 vector is a safe and highly efficient vaccine delivery vehicle. In over one year experiment, none of the vaccinated animals developed adverse effects. We further determined the whole genome sequencing of tissues from one-year vaccinated mice, vector DNA sequences were not detected. Thirdly, rAAV vaccines have been reported to be very stable at −20 degree Celsius or even under room temperature^15^. This indicates that it will be very convenient for storage and transportation, which will largely contribute to the commercialization of the vaccine. Taken together, rAAV5-based RBD-plus vaccine (Covaccinplus) is a very promising candidate vaccine to end this global pandemic.

## Materials and methods

### Viruses and cells

Vero E6 cells (CRL-1586, American Type Culture Collection (ATCC) and HEK293 cells were cultured at 37℃ in Dulbeccos Modified Eagle medium (DMEM) supplemented with 10% fetal bovine serum (FBS), 10 mM HEPES pH 7.4, 1 mM sodium pyruvate, 1X non-essential amino acids, and 100 U/ml of penicillin streptomycin. SARS-CoV-2 strain 2019n-CoV/USA_WA1/2020 and SARS-CoV-2 variants including Alpha, Beta, Gamma and Delta were obtained from Guangdong Provincial Center for Disease Control and Prevention and Institute of Medical Laboratory Animals of Chinese Academy of Medical Sciences. All experiments with infectious SARS-CoV-2 were performed in BSL3 facilities approved Institutional Biosafety Committee.

### Animals in the experiments

6 weeks-old, 36 weeks-old female BALB/c mice and 6-8 weeks female Wistar rats were purchased from the Laboratory Animal Center of Southern Medical University (Guangdong, China). The research was approved by the Guangzhou University of Chinese Medicine Animal Care and Use Committee. All animals were given a commercial mouse food and water ad libitum and housed in a temperature-controlled environment with a 12-hour light-dark cycle.

All animals were immunized with rAAV5-based vaccines or rAAV5-GFP *via* i.m in the hind leg or *via* intranasal inoculation at a single dose (five to eight animals for each group). Sera were collected for cytokines analyses which were performed by the Macau University of Science and Technology using a Non-Human Primate Cytokine Panel kit (MilliporeSigma, Cat# PCYTMG-40K-PX23) on a Bio-Plex 200 instrument (Bio-Rad) according to the manufacturer’s protocol. For virus attacking test, mice were challenged on day 40 after immunization with 3.6Log PFU of SARS-CoV-2 (HRB26M strain) *via* intranasal route. Animals were euthanized and tissues were harvested for further analysis.

### Construction and titration of rAAV5-based vaccines

The recombinant type 5 adenoviral-vectored SARS-CoV-2 vaccines encoding RBD domain (residues 319-541), RBD-plus domain (residues 319-583), S1 protein, full-length of S protein and NTD domain of SARS-CoV-2 were produced by PackGene Biotech (Guangzhou, China). Briefly, rAAV5 packaging plasmids were transfected into HEK293T cells using PEI transfection reagent, according to the manufacturer’s protocol. The transfected cells and supernatants were harvested 72 h post transfection. rAAV5-vaccine was purified and titrated by real-time quantitative PCR (Q-PCR). rAAV5-vaccine was adjusted to 10^12^ GCs/ml in PBS and used for the following vaccinations.

### ELISA

Specific IgG and IgM against COVID-19 in mouse sera were tested by ELISA. Briefly, serially diluted mouse sera were added to 96-well microtiter plates pre-coated with RBD-His or S1-His protein. The plates were incubated at 37□ for 30 min, followed by four washes with PBS containing 0.1% Tween 20 (PBST). Bound Abs were then reacted with HRP-conjugated goat anti-mouse IgG or IgM (Southern Biotech Cat.#1030-05) at 37lJ for 20 min. After four washes, the substrate 3,3,5,5-tetramethylbenzidine (Southern Biotech Cat.#1030-05) was added to the plates and the reaction was stopped by adding 1 N H_2_SO_4_. The absorbance at 450 nm was measured by an ELISA plate reader (Bio-Rad). The endpoint serum dilution was calculated with curve fit analysis of optical density (OD) values for serially diluted sera with a cut-off value of negative control.

### Neutralization assay

Titers of NA in sera of mice immunized with rAAV5 empty or rAAV5-vaccines were detected in Vero E6 cells. Vero E6 cells were seeded at 10^4^/well in 96-well culture plates and cultured at 37°C to form a monolayer. Serial 2-fold dilutions of serum samples were mixed separately with 100 TCID50 (50% tissue-culture infectious dose) of SARS-CoV-2 strain (Guangdong Provincial Center for Disease Control and Prevention and institute of medical laboratory animals of Chinese Academy of Medical Sciences), incubated at 37°C for 1 hour, and added to the monolayer of Vero E6 cells in tetrad. Cells infected with or without 100 TCID_50_ SARS-CoV-2 were applied as positive and negative controls, respectively. The cytopathic effect (CPE) in each well was observed daily and recorded on day 3 post infection. The neutralizing titers of mouse antisera that completely prevented CPE in 50% of the wells were calculated by the Reed-Muench method.

### Quantitative RT-PCR

The viral RNA copies in nasal or lung tissues of challenged mice were determined by quantitative RT-PCR according to the protocol. Total RNA was extracted from 20 mg of lung tissues using a RNeasy Mini kit (company). Then cDNA was synthesized using random primers and the SuperScript II RT kit (company). Extracted RNA (10 μl) was reverse transcribed in a 20-μl reaction mixture containing 1 × first strand buffer, 100 mM DTT, 10 mM each dNTP, 50 ng of random primers, 40 U of RNaseOUT, and 200 U of SuperScript II RT at 42□ for 50 min, followed by 15 min at 70□. The solution was incubated with RNase H (company) at 37□ for 20 min. Synthesized cDNA was quantified using Power SYBR Green PCR Master Mix (company) in a 20-μl mixture containing 5 μl of cDNA (1/10), 10 μl of 2 × Power SYBR Green PCR Master Mix, 3 μl of RNase-free H_2_O, 10 μM forward primer and reverse primer in a Mx3000 QPCR System (company).

### Cell surface markers/intracellular cytokines staining

Single-cell suspensions (3×10^6^) from spleens of the vaccinated mice were harvested and stimulated with or without SARS-CoV-2 S-specific peptide (1μg/ml) plus anti-mouse IL-2 (20 U/ml). Cells with stimulatory agents were incubated for 72 hour at 37□ with 5% CO_2_. The cells were harvested and stained directly with conjugated mAbs specific for cell surface markers including CD45, CD3, CD4, CD8, CD44, CD62L for 30 min at 4°C. Intracellular antigens including IFN-γ, IL-4 and TNF-α were stained for 20 minutes in the dark at room temperature. The stained cells were analyzed using a flow cytometer (BD company). Data were analyzed by Cell Quest software (company).

### Statistical analysis

Values were presented as mean with SEM. Statistical significance among different vaccination groups was calculated by the student t test using Stata statistical software (GraphPad Prism 7). Values of P<0.05 were considered significant.

## Acknowledgements

This study was partially funded by Guangzhou Laboratory (Grant No. EKPG21-23), the Science Research Project of the Guangdong Province (Grant No. 2020A1111340003) and China Postdoctoral Science Foundation (Grant No. 2020T130029ZX). Guangdong Hengda Biomedical Technology Co., Ltd. (Guangzhou, China) provided RBD-plus vaccines for the studies. The PackGene Biotech Co., Ltd. (Guangzhou, China) provided supports in production of RBD-plus vaccines. The GMP level plasmids were produced by Yaohai Bioharmaceutical Co., Ltd. (Taizhou, China). Whole Genome Sequencing technology was provided by BGI Genomics Co., Ltd. (Guangzhou, China). SARS-CoV-2 virus challenge tests were carried out in Harbin Veterinary Research Institute, Harbin, China.

